# Multistage activity within a diverse set of epi-drugs against *Plasmodium falciparum* parasites

**DOI:** 10.1101/692913

**Authors:** Nanika Coetzee, Hilde von Grüning, Mariette van der Watt, Janette Reader, Lyn-Marié Birkholtz

## Abstract

The epigenome of the malaria parasite, *Plasmodium falciparum*, is associated with control of various essential processes in the parasite including control of proliferation of asexual development as well as sexual differentiation. The unusual nature of the epigenome has prompted investigations of the potential to target epigenetic modulators with novel chemotypes. Here, we explored the diversity associated with a library of 95 compounds, active against various epigenetic modifiers within cancerous cells, for activity against multiple stages of *P. falciparum* development. We show that *P. falciparum* is differentially susceptible to epigenetic perturbation during asexual and sexual development, with early stage gametocytes particularly sensitive to epi-drugs targeting both histone and non-histone epigenetic modifiers. Moreover, 4 compounds targeting histone acetylation and methylation, show potent multistage activity against asexual parasites, early and late stage gametocytes, with transmission-blocking potential. Overall, these results warrant further examination of the potential antimalarial properties of these hit compounds.

## Background

The almost inevitable nature of drug resistance development by malaria parasites enforces continued exploration of novel classes of antimalarial drugs. To contribute to global malaria elimination strategies (1, 2), such compounds would need to target multiple life cycle stages including the rapidly dividing asexual parasites for chemotherapies, as well as terminally differentiated sexual gametocytes for transmission-blocking activity. Importantly, to prolong resistance development, new chemical matter targeting novel biological activities in the parasite is needed.

In oncology research, epigenetic therapeutics (‘epi-drugs’) that inhibit epigenetic modulators evidently hold great promise as targets for anticancer therapies (3). The result is an epigenetic de-regulation with antitumor activity, due to targeting histone modifying enzymes and DNA methyltransferases, or facilitating reader interference by inhibiting or activating epigenetic processes. This results in particularly histone post-translational modifications (PTMs) and disruption of transcriptional processes, chromatin structure maintenance and DNA repair (4, 5). Various anticancer epi-drugs have been approved for clinical use, including Azacitidine, Decitabine, Vorinostat and Romidepsin (6).

*Plasmodium falciparum* relies heavily on epigenetic mechanisms to drive both asexual proliferation and sexual differentiation (reviewed in (7–11)). The parasite’s genome encodes a unique complement of histone modifying enzymes including histone deacetylases (HDACs), histone acetyltransferases (HATs), histone methyltransferases (HMTs, including lysine HKMT), protein arginine methyltransferases (PRMTs), and histone demethylases (HDMs) (11) in addition to other non-histone epigenetic modifiers. As a result, inhibitors of particularly histone modifying enzymes have been investigated as novel chemotypes in antimalarial drug discovery efforts (12–22).

Selective anticancer epi-drugs have been investigated for their activity against *P. falciparum* asexual parasites and to a lesser extent, against gametocyte stages. These compounds disturb the parasite’s gene expression, ultimately leading to cell death (20–22). HDACs are seen as particularly promising drug targets due to resultant hyperacetylation (on various histone sites) upon disruption of these activities, with HDACi (HDAC inhibitors) receiving the vast majority of interest. This includes well-known hydroxymate-based inhibitors like SAHA (suberoylanilide hydroxamic acid, Vorinostat and its derivates) and TSA (Trichostatin A) as well as cyclic tetrapeptides like apicidin, which have shown selective inhibition against asexual *P. falciparum* stages (17, 22–26) and gametocytes (17, 26). SAHA additionally retained activity in clinical isolates of both *P. falciparum* and *P. vivax* (27). These data have led to larger screens for diverse and selective inhibitors of HDACs (14, 28–30). In *P. falciparum*, histone lysine methyltransferases (HKMTs) are involved in both transcriptional activation (through H3K4me marking) and repression (e.g. H3K9me marks), and are also hypothesised to be promising drug targets (16), with BIX01294 (as model HKMT inhibitor, HKMTi) successfully inhibiting asexual *P. falciparum* proliferation and gametocyte viability (15, 16). The diaminoquinazoline chemotype has shown to be particularly effective HKMTi against asexual *P. falciparum* parasites, with diversity set screens resulting in selective inhibitors identified (15, 16, 19). Although these data support the notion that epigenetic modulators could be drug targets in parasite development as well as differentiation, some chemotypes show overt toxicity, poor selectivity and sometimes poor pharmacokinetics (31). Diverse chemotypes targeting various epigenetic modulators should therefore be explored.

In this study, a library of anticancer compounds (Cayman Epigenetics Screening Library, Cayman’s Chemicals, USA) with known capabilities to inhibit diverse epigenetic modulators in cancerous cells, was evaluated for their antiplasmodial activity against multiple *P. falciparum* stages. The library consists of 39% HDACi and 15% HKMTi; with the remaining compounds divided into 11 other inhibitor subtypes including targeting of HAT, DNA demethylases (DNDM), DNA methyltransferases (DNMT), protein arginine deiminases, PRMT, bromodomain proteins, HDMs, lysine-specific demethylases (LSD), and processes involved in hydroxylation and phosphorylation. As the unusual epigenome and associated regulatory machinery of the parasite provide extensive biology to be investigated, the use of this diverse library of epi-drugs could prioritise which epigenetic modifiers have potential as novel druggable entities. This study describes a comprehensive screening of inhibitors of epigenetic modulators against multiple life cycle stages of *P. falciparum*, including asexual parasites, early (immature) and mature late stage gametocytes and gamete formation. We demonstrate that HDACi and HKMTi remains the most potent compounds with multistage activity but identify new chemotypes with the potential to be used as chemical starting points for antimalarial drug discovery efforts.

## Results

### Comparative profiling of the Cayman Epigenetics library for inhibition activity against *P. falciparum* parasites

All 95 compounds in the Cayman Epigenetics library were profiled for *in vitro* activity against asexual and sexual *P. falciparum* parasites in a dual-concentration screen (1 and 5 µM, Figure 1A, supplemental figure 1, supplementary file 1). This included stage-specific evaluation of the compounds against early (>85% stage II/III) and late stage (>95% stage IV/V) gametocytes. The majority of the compounds (76% against asexual parasites, early (69%) and late (82%) stage gametocytes) showed no/minimal activity. Although similar hit rates and compound identities were observed between asexual parasites and early stage gametocytes (24 and 30% of compounds, respectively, active against these stages at >50% inhibition, Pearson correlation r^2^ of 0.5), the distribution of compounds displaying moderate activity against early stage gametocytes were almost double that against asexual parasites (18 *vs.* 10%). Overall, the compounds were the least active against late stage gametocytes (18% hit rate). Additionally, the nature of the compounds active against asexual parasites and late gametocytes (and between those active against early and late stage gametocytes) showed poor correlation (r^2^ of 0.3 and 0.2, respectively), indicating some stage-specificity in the distribution of the compounds active against each stage.

**Figure 1.**
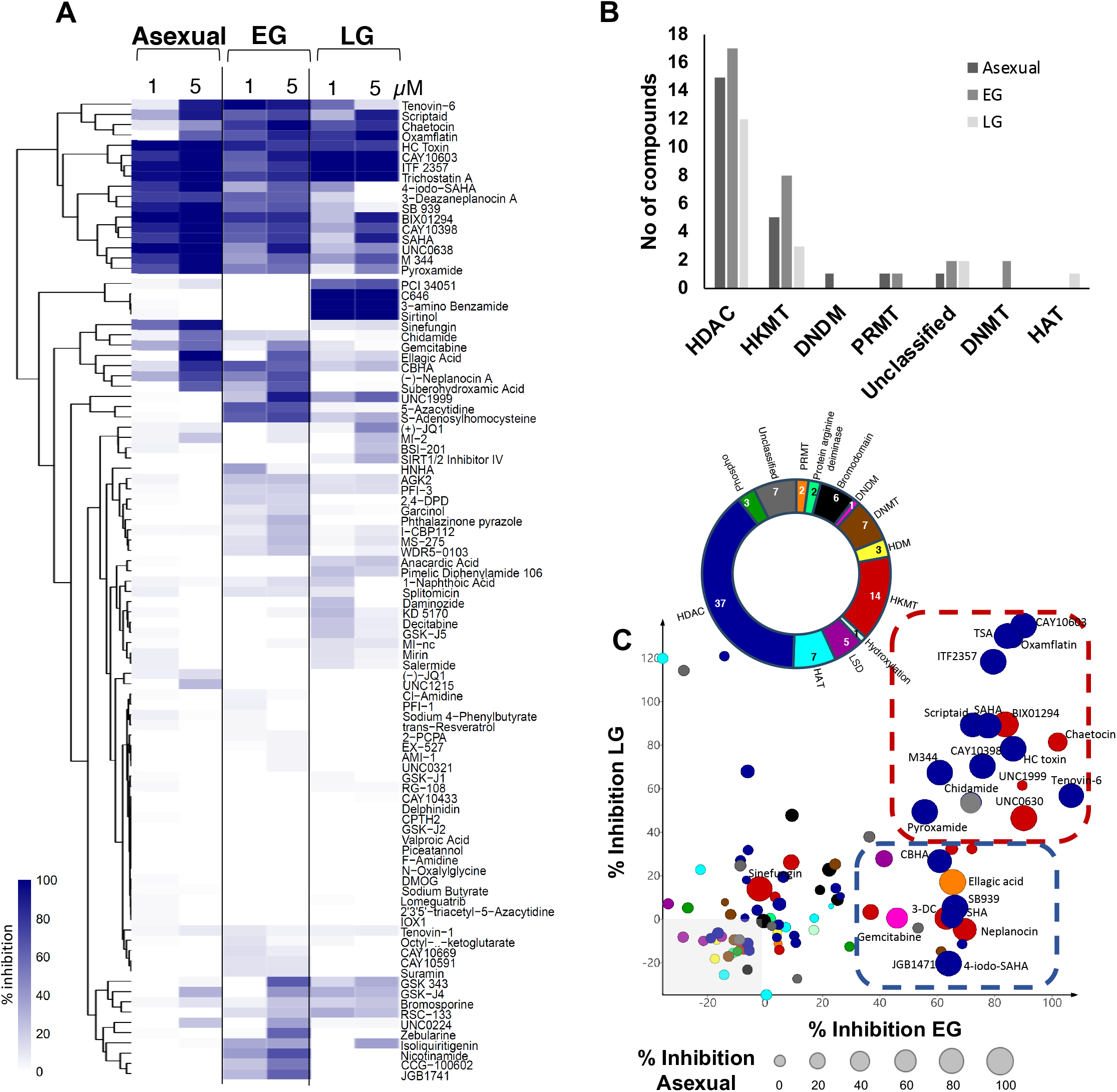
Comparative profiling of the Cayman Epigenetics library of drugs screened for inhibition activity against *P. falciparum* parasites. **(A)** Primary compound screening of 95 drugs that inhibit epigenetic modulators was performed using the SYBR Green I-based fluorescence assay for asexual parasites (strains 3D7, 96 h drug pressure on ring stage parasites) and the pLDH assay for early and late stage gametocytes (strain NF54, 72 h drug pressure each). The heatmap shows inhibition of asexual parasites and early (EG) and late stage (LG) gametocytes at 1 and 5 µM drug pressure. The color scale indicates percentage activity of the treated control after background subtraction of the positive controls (chloroquine for asexual parasites and methylene blue for gametocytes). Compounds with similar inhibition profiles were hierarchically clustered based on Euclidean distance using R Software (v3.6.0). **(B)** Distribution of compounds with >50% activity per life cycle stage based on their inhibitor classification within the Caymans library. **(C)** Epi-drug library composition based on inhibitor classification, targeting epigenetic modifiers, with the number of compounds per class indicated. Protein arginine methyltransferase (PRMT), DNA demethylase (DNDM), DNA methyltransferase (DNMT), histone demethylase (HDM), histone lysine methyltransferase (HKMT), lysine-specific demethylase (LSD), histone acetyltransferase (HAT), histone deacetylase (HDAC). Inhibition at 5 μM (%) was compared between asexual parasites (circle size; n=3) and early (EG) & late (LG) stage gametocytes (n=1); separated based on the inhibitor type (colour scale corresponding to inhibitor classification as in B). Compounds with multi-stage activity is identified in the red block and those with asexual and EG preference in the blue block. SHA: suberohydroxamic acid; 3-DC: 3-deazaneplanocin.

Hierarchical clustering of the compounds based on Euclidean distances further revealed this stage-specific distribution (Figure 1A). A subset of 17 compounds (including well known compounds like TSA, SAHA, BIX01294 and HC Toxin), display activity against all life cycle stages tested, marking these compounds as multistage inhibitors. However, 4 compounds (PCI34051, C646, 3-amino benzamide and sirtinol) show late stage gametocyte preference with an additional 10 compounds clustered due to increased activity towards early stage gametocytes.

The multistage activity of the 95 compounds was stratified based on the inhibitor classes descriptors for these compounds (Figure 1B). As expected, the majority of the active compounds are classified as HDACi and HKMTi, and compounds from these classes target all life cycle stages (Figure 1C). Interestingly, compounds classified in the library as potential DNDM, DNMT, PRMT and HAT inhibitors were within the ‘hit’ pool. However, inhibitors targeting bromodomain proteins, hydroxylation, phosphorylation, and demethylation activities (both histone demethylase and lysine-specific demethylase) were not particularly active (<50% inhibition). Compounds with >50% inhibition (at 5 μM) against all three parasite stages included mostly HDACi (Scriptaid, HC Toxin, ITF 2357, Tenovin-6, CAY10603, M 344, Oxamflatin, Pyroxamide, Trichostatin A, CAY10398, SAHA, Chidamide) and some HKMTi (Chaetocin, UNC0638, BIX01294) (Figure 2C). Within these compounds, some showed distinct dual stage-specific activity against asexual and early gametocyte stages (CBHA, Ellagic Acid, SB939, Suberohydroxamic acid, 3-Deazaneplanocin A, (-)-Neplanocin A, 4-iodo-SAHA), or both gametocyte stages (UNC1999) (Figure 1C). An additional subset of compounds is solely active against asexual stages and early stage gametocytes and include DNDMi (gemcitabine), PRMTi (ellagic acid). Stage-selective activity was identified for sinefungin, targeting only asexual parasites or early stage gametocytes (S-Adenosylhomocysteine, 5-Azacytidine, GSK 343, Nicotinamide, JGB1741, Zebularine, CCG-100602) or late stage gametocytes (3-amino Benzamide, C646, Sirtinol, PCI 34051) being targeted.

**Figure 2.**
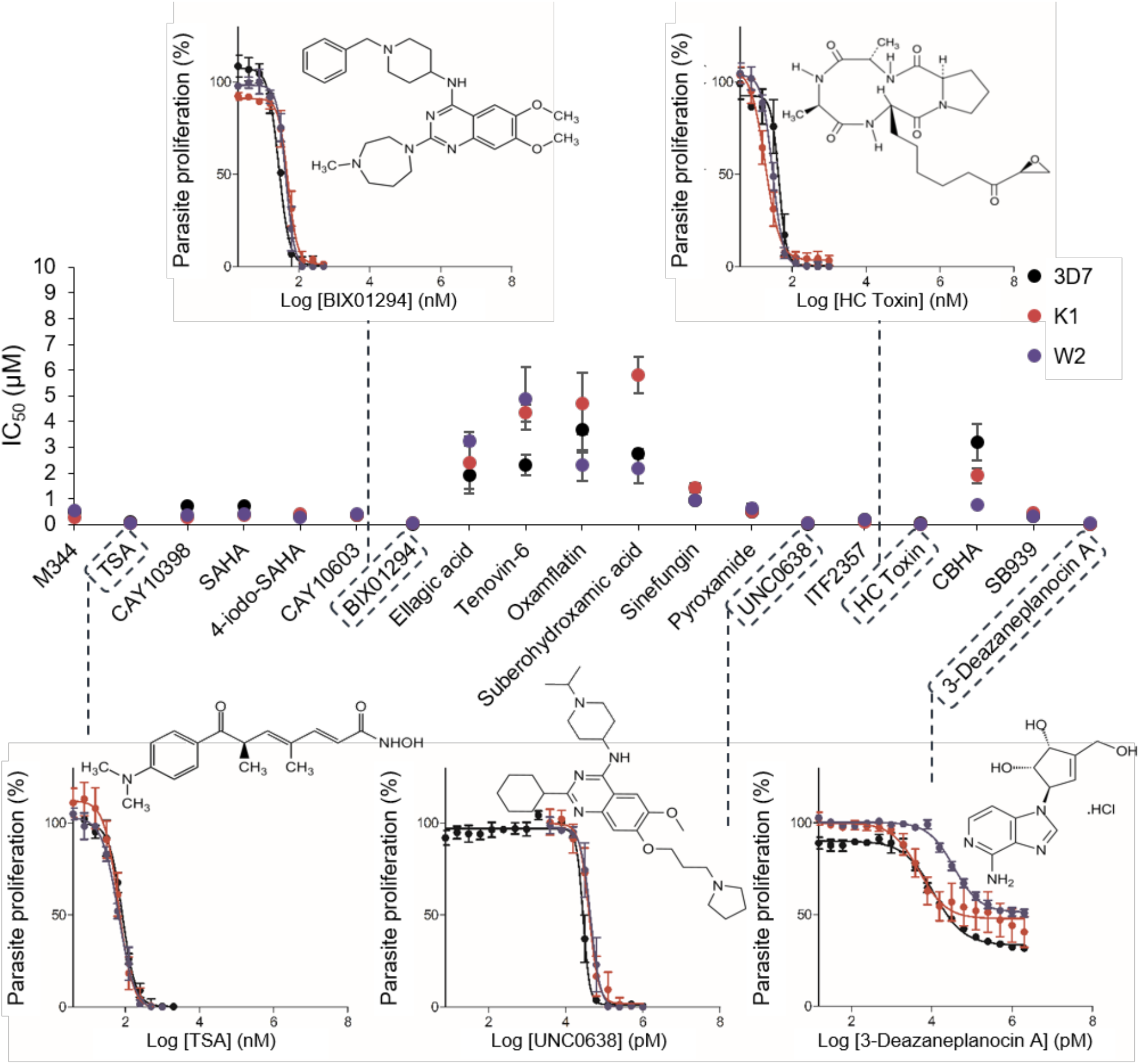
*In vitro* activity of the most active epi-drugs against asexual drug sensitive and resistant *P. falciparum* strains. Compounds were screened using the SYBR Green I-based fluorescence assay to determine dose-response against 3D7 (drug sensitive, black), K1 (drug resistant, red) and W2 (drug resistant, blue). Data are represented as a percentage of untreated control to determine cell proliferation. Sigmoidal dose-response curves were plotted using GraphPad 5.0, from which the IC_50_ values could be determined. Results for all compounds are representative of three independent biological replicates (n=3 ± SEM).

### Antiproliferative activity against asexual parasites

Of the hits described above, 19 compounds were selected for IC_50_ determination against both drug sensitive (3D7) and resistant (K1 and W2) strains of *P. falciparum* parasites (Figure 2A, Table 1). Collectively, IC_50_ values ranged between 7 nM to 6 μM for all the parasite strains evaluated. The resistance indices (RI; ratio of the IC_50_ value of the resistant strain to the sensitive strain, i.e. K1/3D7 and W2/3D7) averaged at 1.12, indicating limited cross-resistance to the K1 and W2 *P. falciparum* parasite strains (Table 1). Five compounds were extremely potent at <100 nM against all parasite strains, with the most active compounds (3-Deazaneplanocin A, BIX01294, UNC0638) classified as HKMTi, followed by two HDACi, HC Toxin and TSA. TSA, BIX01294 and SAHA showed similar IC_50_ values to those found in previous reports (16, 17, 22). 3-Deazaneplanocin A could, however, consistently not cause complete parasite clearance, even up to 100 μM or prolonged exposure (up to 96 h tested). All of the compounds with activity below 500 nM belong to the HDACi class, emphasizing the importance of this activity for asexual proliferation of *P. falciparum*.

**Table 1:**
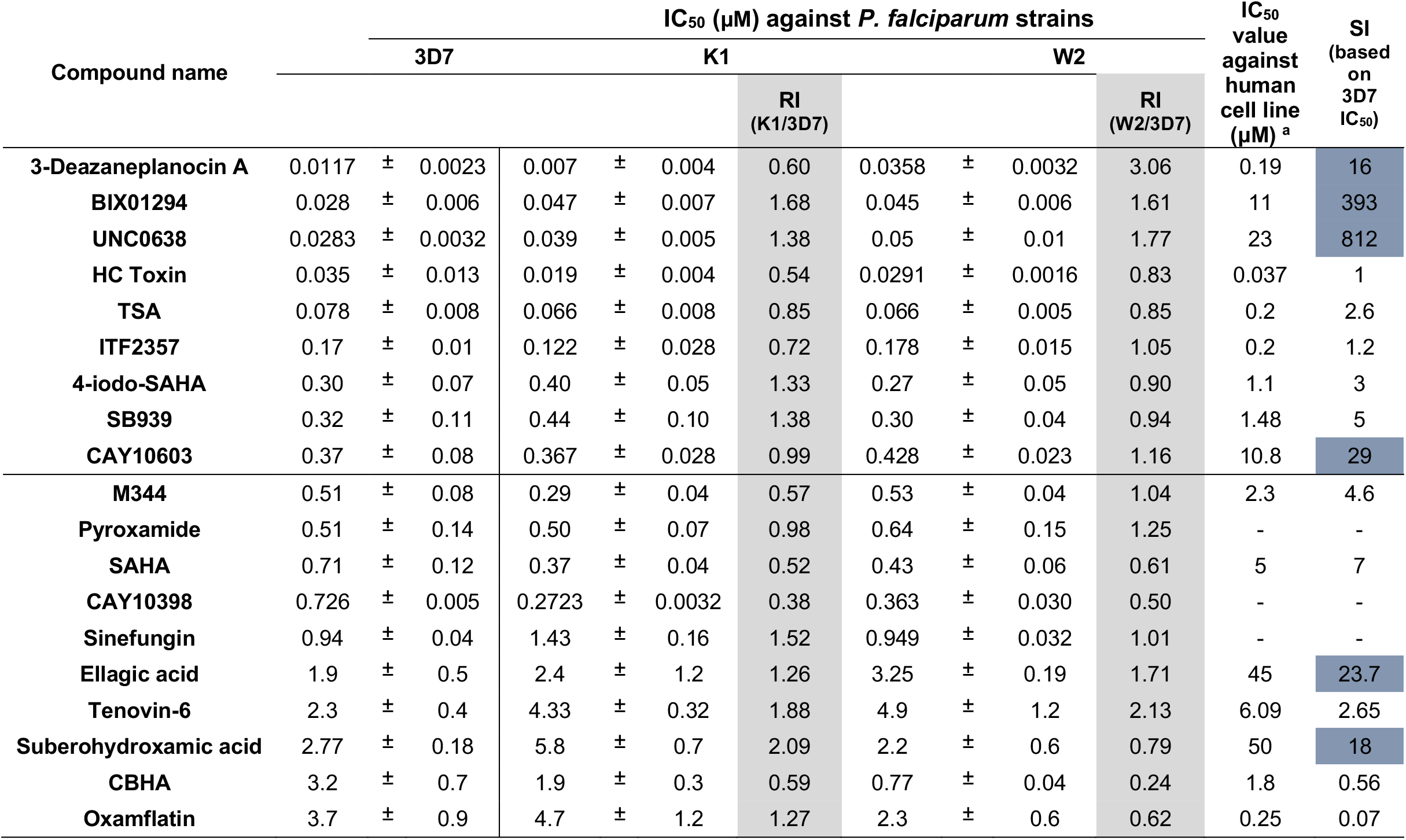
Cross-resistant dose-response for the active epi-drugs. Compounds were screened for IC_50_ against 3D7, K1 and W2 asexual parasite strains using the SYBR Green I-based fluorescence assay. The resistance index (RI; shaded grey) and selectivity index (SI) for each compound is shown. Compounds with SI˃10 are shaded in blue. Activity data of the compounds on various mammalian lines were collated from previous reports (15, 23, 43, 55-59). Results for all compounds are representative of three independent biological replicates with technical triplicates (n=3, IC_50_ ± SEM).

Since the compounds in the library were included based on evidence of activity against mammalian lines, their selective toxicity towards *P. falciparum* parasites was determined (Table 1). All of the most potent compounds, except for HC Toxin, showed preference towards *P. falciparum* parasites, with particularly BIX01294 and UNC0638 highly selective towards the parasite with SI>300, supporting that HKMT activity is essentially important to malaria parasite proliferation. CAY10603 and suberohydroxamic acid (HDACi) and ellagic acid (a PRMTi), were also 10-fold more active against the parasite than mammalian cells with SI values >10. These compounds therefore result in a base set of chemical entities that can be explored in medicinal chemistry programs, with selective inhibition towards malaria parasites.

### Gametocytocidal and gametocidal activity

Selected compounds were evaluated for their activity on both early and late stage gametocytes, with 8 compounds showing activity at <5 μM on gametocytes (Figure 3). Comparative low μM activity was observed against both early and late stage gametocytes for the HKMTi Chaetocin (IC_50_: 0.92 ± 0.29 μM early gametocyte; 1.34 ± 0.17 μM late gametocyte) and for 5 HDACi:CAY10603 (IC_50_: 1.6 ± 0.8 μM vs 1.30 ± 0.29 μM early vs late gametocyte), ITF 2357 (IC_50_: 3.0 ± 0.9 μM early gametocyte IC_50_; 2.23 ± 0.09 μM late gametocyte), Oxamflatin (1.2 ± 0.6 μM early gametocyte IC_50_; 3.0 ± 0.4 μM late gametocyte IC_50_), HC Toxin (2.817 ± 0.269 μM early gametocyte IC_50_; 2.25 ± 0.54 μM late gametocyte IC_50_). The HDACi Scriptaid had marginal preference towards late stage gametocytes (3.16 ± 0.625 μM early gametocyte IC_50_; 1.096 ± 0.02 μM late gametocyte IC_50_), similarly to sirtinol with potent activity against these more mature gametocytes (0.112 ± 0.015 μM late gametocyte IC_50_). Activities for known compounds SAHA, BIX01294 and TSA were comparative to reported values (SAHA: (1.41 ± 0.13 μM early gametocyte IC_50_; 0.81 ± 0.21 μM late gametocyte IC_50_ (17); BIX01294: 0.014 ± 0.002 μM early gametocyte IC_50_; 5.86 μM late gametocyte IC_50_ (15); TSA: 0.09 ± 0.01 μM early gametocyte IC_50_; 4.67 ± 1.09 μM late gametocyte IC_50_ (17)).

**Figure 3.**
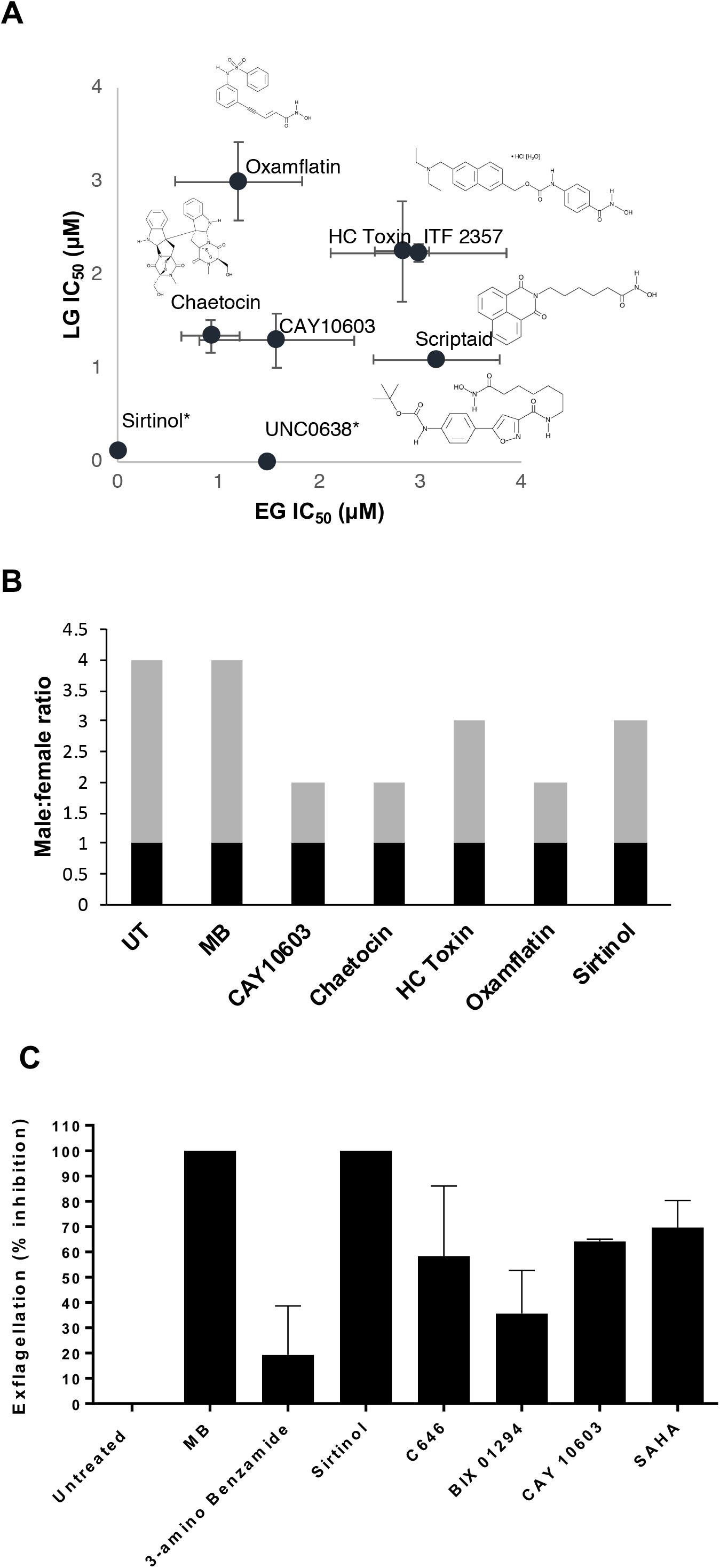
Gametocytocidal activity of the selected epi-drugs against early and late stage gametocytes of *P. falciparum*. **(A)** Compounds were screened using the pLDH assay to determine dose-response against early (>85% stage II and III, EG) and late (>95% stage IV/V, LG) gametocytes after drug pressure for 72 h. Data are represented as a percentage of untreated control. Sigmoidal dose-response curves were plotted using GraphPad 5.0, from which the IC_50_ values could be determined. Results for all compounds are representative of three independent biological replicates (n=3 ± SEM), except for sirtinol, scriptaid, HC toxin and UNC0638 (n=1). Where not show, error bars fall within the symbol. Sirtinol was not tested against EG and UNC0638 was not tested against LG. **(B)** Male:Female gametocyte ratio (males in black bars, females in grey bars) affected by selected compounds. Gametocytes were binned morphologically after evaluating >1000 erythrocytes each on Giemsa stained slides. **(C)**The ability of selected compounds to inhibit male gamete formation. The inhibition of exflagellation of male gametes (% exflagellating males compared to untreated controls) was visually assessed for two independent biological repeats from 15 videos per repeat, ± SEM.

CAY10603, Chaetocin and Oxflamfatin was able to reduce the normal ~3:1 ratio of female:male mature gametocytes to equal proportions (Figure 3B). In addition, Chaetocin and sirtinol, and to a lesser extent CAY10603, affected the functional viability of male gametocytes to exflagellate (>60% inhibition) (Figure 3C). With these also affecting female gametocytes, it points to shared biological activities being targeted rather than sex-specific processes.

### HDACi and HKMTi as multistage targeting epi-drugs

As the HDACi and HKMTi showed the most extensive activity against multiple *P. falciparum* parasite stages, the potential for structure activity relationships between these two inhibitor classes were explored (Figure 4). Although the majority (>80%) of the compounds were not structurally related, even though they did show activity in the primary screen against multiple stages, a core hydroxamate-based scaffold could be identified for 5 HDACi, which showed ˃80% structural similarity, including Pyroxamide, SAHA, 4-iodo-SAHA, CAY10433 and Pimelic Diphenylamide 106. Conversely, 4 HKMTi showed ˃80% structural similarity as 4-quinazolinamine-based structures, including BIX01294, UNC0638, UNC0224 and UNC0321. A few other structurally similar pairs were also identified, including CAY10398 and M 344 (both HDACi) that has multistage activity and only differs with a single backbone carbon. These compounds show promise as chemical starting points for antimalarial optimisation as they have unique chemical scaffolds compared to most compounds and have an altered molecular target.

**Figure 4.**
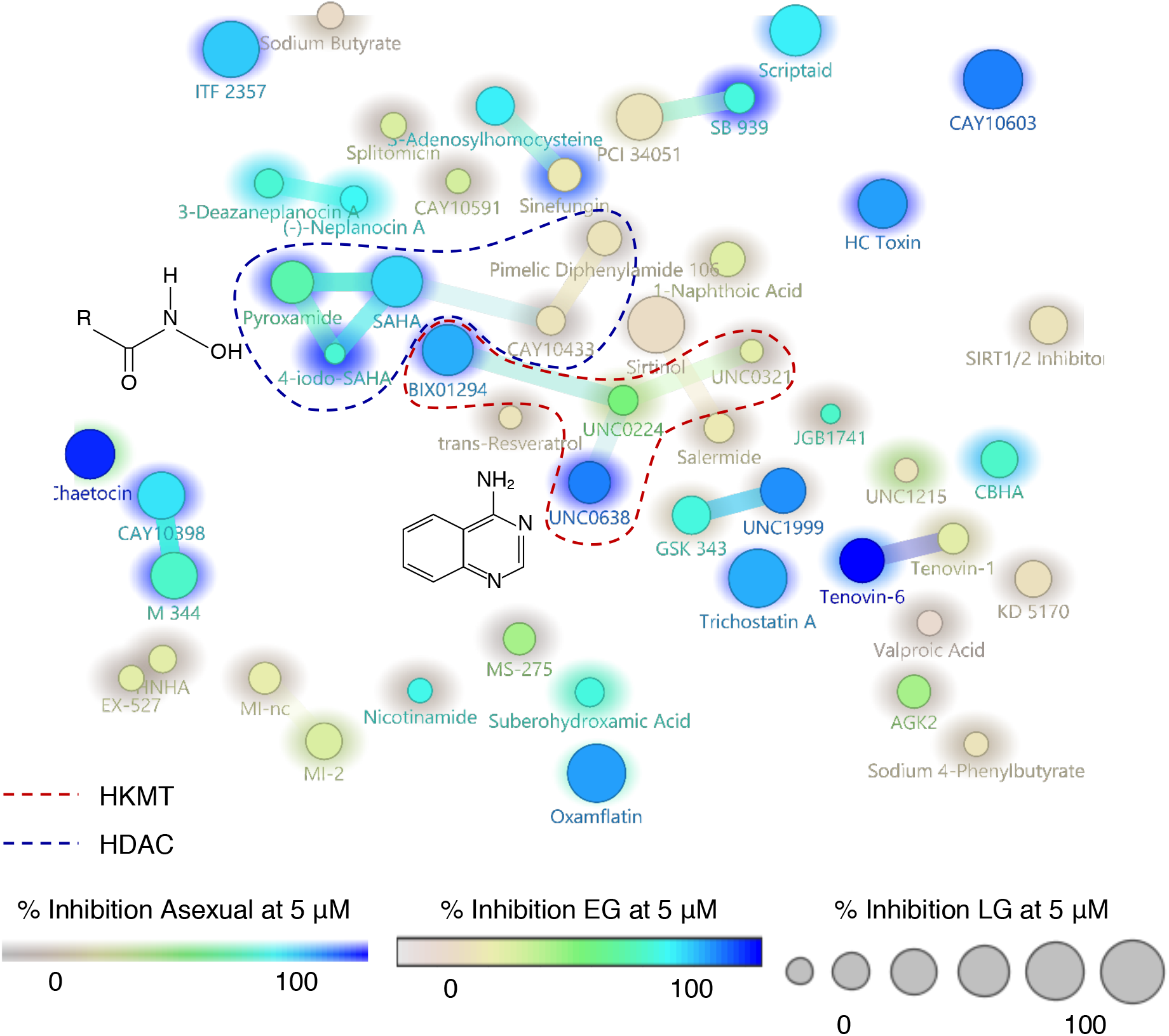
Structure activity relationship within the HDAC and HKMT inhibitor series. Structural feature (SkelSphere) analysis was performed with superimposed activity cliff analysis (Osiris DataWarrior V4.2.7) at 80% structural similarity cut-off. Compounds are limited to HDAC (blue line) and HKMT (red line) inhibitors. Inhibition at 5 μM (%) was compared between asexual parasites (background shading; n=3), early (EG; circle colour; n=1) and late (LG; circle size; n=1) stage gametocytes. The active chemical groups shared between two major inhibitor groups are shown; hydroxamate for HDAC and 4-quinazolinamine for HKMT.

## Discussion

The *P. falciparum* parasite’s epigenetic regulatory machinery has previously been shown to be a valuable drug target given the importance of gene expression for parasite development, but remains to be exploited in full (15, 16, 22, 23, 32-35). In this study, various compounds, known to target the epigenetic modulators, were investigated for their antiplasmodial activity against asexual parasites, early and late stage gametocytes and gametes. Potent chemical scaffolds were identified with multistage activity, and this study again highlights the potential of HDACi and HKMTi to be exploited as lead compounds upon further experimental validation, cytotoxicity reduction and chemical property optimisation.

The majority of the epigenetic inhibitor classes represented in the library did not show any activity against any of the three *P. falciparum* stages tested. Barring compound uptake issues, this suggests that the epigenetic modulators that should have been targeted by compounds in the library are 1) not essential in the survival of the parasite through development, 2) that the homologues for these epi-drugs’ mammalian targets are not found in *P. falciparum* or 3) cannot be targeted with the compounds within this library. These epigenetic inhibitor classes included phosphorylation, protein arginine deiminase, HDM, hydroxylation, LSD and bromodomain inhibitors. Amongst the non-histone epigenetic modulators that were targeted, DNDMi, DNMTi and PRMTi were identified. DNMTi were recently identified with *in vitro* activity against *P. falciparum* and retains oral efficacy in a *P. berghei* mouse model (14). These compounds are selective inhibitors of human DNMT3a (14), with activity only shown against asexual *P. falciparum* parasites. These DNMTi does not bear any structural relationship with the compounds tested from the Caymans library, where only Zebularine was identified as DNMTi, with selective inhibition against early gametocytes. The DNDMi Gemcitabine was also shown to have stage-specificity for asexual stages, suggesting possible and antagonistic involvement of DNA methylation-linked chromatin condensation in these stages of development (36). PRMTi were moderately active against asexual and early gametocyte stages. This includes Ellagic Acid, which was previously indicated as potent against asexual parasites (37). Although antioxidant pleiotropic effects of this compound cannot be ignored, the multistage activity reported here suggests a possible role of arginine methylation during these stages (38), but not in mature gametocytes.

Histone-associated epigenetic regulators remain overrepresented as targets, both due to the composition of the library screened, but also due to the known importance of histone acetylation and methylation to parasite survival (21, 39). However, HDACs and HKMTs remain the activities targeted the most frequently; a single HATi was identified with multistage activity (C 646, specific competitive inhibitor of human p300 HAT). Hydroxamate-based HDACi and 4-quinazolinamine-based HKMTi remained the most potent chemical scaffolds that target *Plasmodium* parasite development at the symptomatic asexual and transmissible gametocyte forms. Hydroxamate-based HDACi were previously shown to have potent antiplasmodial activity with limited cytotoxicity, and this class contains some clinically approved compounds which have been attempted in drug repurposing studies (22, 24, 40-43). These compounds lead to DNA hyper-acetylation, resulting in the de-regulation of transcription and ultimately cell cycle arrest and cell death (25, 26, 44, 45). Some HDACi were also completely pan-inactive, thereby suggesting that the parasite relies on a specific set of HDACs to regulate its chromosomal condensation via acetylation (reviewed in (46)). The hit HDACi selected in this study display unique chemical scaffolds and showed activity against all stages, although not at equipotent levels (CAY10603, ITF 2357 and Oxamflatin), with potenty increased against asexual parasites and early stage gametocytes. Comparatively, HKMTi overall has the best potency and selectivity with additional activity retained against transmissible stages, similar to previous reports on 4-Quinazolinamine-based HKMTi (15, 16). Selective inhibition of HKMT activity can either lead to an increase or decrease in transcription, depending on the position and degree of methylation and ultimately contributes to transcriptional de-regulation and cell death (16). For instance, the HKMTi, 3-Deazaneplanocin A, selectively inhibits H3K27me3 and H4K20me3, and reactivates silenced developmental genes in cancer cells that are not silenced by DNA methylation (47).

The stage-specific inhibition profiles observed for the wide variety of epi-drug inhibitor classes support the findings that the parasite makes use of altered epigenetic regulatory mechanisms to differentiate itself during asexual proliferation and sexual differentiation (7, 48, 49). Selective *Plasmodium* inhibition was only shown for 6 compounds of the series, which suggests that the epigenetic modulators targeted by these compounds (HKMT, HDAC and PRMT) show diversity between the parasite and human homologues, as previously shown by the unique set of *Plasmodium*-specific epigenetic factors that differs vastly from those in its mammalian host (39). The primary evaluation of the activity of epi-drugs against the multiple life cycle stages indicated that early stage gametocytes were particularly susceptible to epi-drug inhibition, supported by reports of a unique epigenetic repertoire associated with these stages, where the switch between asexual and sexual stages was accompanied by dynamic histone PTM landscape alterations (7). The differentiation between the compounds active against the different life cycle stages correlates with unique and stage-specific histone PTM dynamics during the parasite’s life cycle, with clear peaks abundances for some epigenetic marks associated with particular life cycle stages (7, 48, 49) and highlights the importance of these activities to parasite development.

Collectively, the data implies that the epigenetic modulators affected are essential for parasite development throughout its asexual and sexual life cycle. Our study reveals that certain chemical scaffolds shared between multistage active compounds hold potential as chemical starting points for further development of derivatives with increased potency, selectivity, or improved physico-chemical properties.

## Materials & Methods

### Asexual *P. falciparum* parasite cultivation and antiproliferative assays

*P. falciparum* parasites were maintained at 37°C in human erythrocytes suspended in complete culture medium and ring-stage synchronised as described (50). SYBR Green I fluorescence was used to determine compound activity against asexual ring stages (1% haematocrit, 1% parasitaemia), treated with compounds at 1 and 5 μM (for primary screening of inhibitory activity), for 96 h at 37°C as described (50, 51), chloroquine disulphate (1 µM) as positive drug control. SYBR Green I fluorescence was measured using a Fluoroskan Ascent FL microplate fluorometer (Thermo Scientific, excitation at 485 nm and emission at 538 nm). Unless otherwise stated, each compound was screened in technical triplicates for at least three independent biological replicates (n=3). Hit compounds were selected for full dose-response evaluation under the same assay conditions as above, but against drug sensitive (3D7) and drug resistant W2 (chloroquine, quinine, pyrimethamine and cycloguanil resistant) and K1 (chloroquine, pyrimethamine, mefloquine and cycloguanil resistant) *P. falciparum* strains to determine inhibitory concentrations of the compounds needed to affect 50% of the parasite population (IC_50_). Data are representative of at least 3 biological repeats, performed in technical triplicates. Assay performances were evaluated with average %CV at 4.47 and Z-factors at >0.6.

### Parasite lactate dehydrogenase assay to determine inhibition against the early and late stage *P. falciparum* gametocytes

Gametocytes were induced from asexual *P. falciparum* NF54 parasites as described before (50). The parasite lactate dehydrogenase (pLDH) assay was performed as previously described (50, 52, 53). Early and late stage *P. falciparum* gametocyte cultures (2% haematocrit, 5% gametocytaemia) were treated with compounds at 1 and 5 μM (primary dual-point primary screening), with methylene blue (5 µM) as positive control for inhibition. Gametocytes were treated under drug pressure for 72 h at 37°C, followed by drug washout with culture medium and remaining pLDH activity was determined 72 h later by addition of Malstat reagent (53) and absorbance measured with a Multiskan Ascent 354 multiplate scanner (Thermo Labsystems, Finland) at 620 nm. A full dose-response evaluation was performed for hit compounds against early and late stage *P. falciparum* gametocytes for three independent biological replicates. Assay performances were evaluated with average %CV at 4.54 and Z-factors at >0.5.

### Data analysis

Chemi-informatics evaluation of compound activities including structure-activity landscape analysis was performed with Osiris DataWarrier v4.7.2

### Gamete formation assays

Male and female gametocytes were detected visually with Giemsa stain after 48 h drug pressure, by counting at least 1000 cells. The male exflagellation inhibition assay (EIA) was performed by capturing movement of exflagellation centres over time by video microscopy (adapted from Ghosh *et al.*, 2010). Mature gametocyte culture (>95% stage V; 1 ml) was centrifuged at 3500 rpm for 30 seconds and the pellet resuspended in 30 µl of ookinete medium (RPMI-1640 medium containing L-glutamine (SIGMA, R6504), 0.024 mg/ml gentamycin (HyClone, SV30080.01), 202 µM hypoxanthine (SIGMA, H9636), 25 mM HEPES (SIGMA, H4034), 0.2% Glucose (SIGMA, G6152), 0.5% (w/v) Albumax II (Invitrogen, Paisley, UK) and supplemented with 20% human serum). The activated culture (10 µl) was introduced into a Neubauer chamber and placed on the microscope platform to settle homogenously. Time was noted as time zero (T_0_) and the chamber incubated at room temperature (RT). Movement was recorded by video microscopy using a Carl *Zeiss NT 6V/10W Stab* microscope, fitted with a MicroCapture camera at 10X magnification and then quantified by a semi-automated method, a modification of the method described by (54). A series of 15 videos of 8-10 seconds each were captured at random locations between minute 15 and 22.5 after incubation. Each video was analysed using Icy bio-image analysis software in order to quantify the number of exflagellating centres.

## Author Contributions

LMB conceived the study. NC, HvG, MvdW, JR conducted experiments, all authors interpreted results. NC, LMB wrote the paper with inputs from the other authors. All co-authors approved the final version of the paper.

## Acknowledgements

The UP ISMC acknowledges the South African Medical Research Council (SA MRC) as Collaborating Centre for Malaria Research. This work was supported by the South African Research Chairs Initiative of the Department of Science and Technology, administered through the South African National Research Foundation (UID 84627) to LMB.

