## Supplementary figures and images for "Multistage activity within a diverse set of epi-drugs against *Plasmodium falciparum* parasites"

### Supplemental figure 1

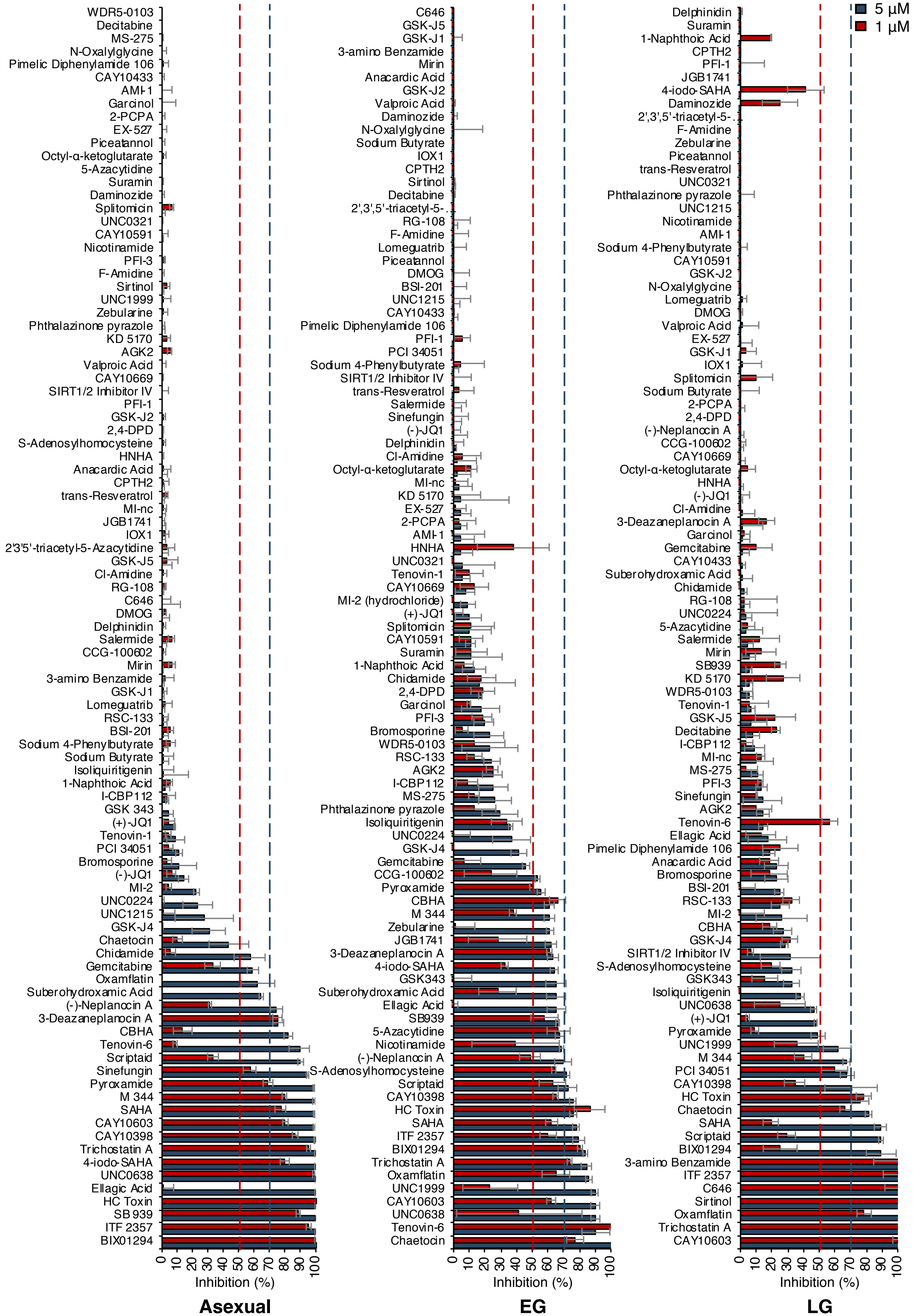
